# OptiRanker: an Advanced Tool for Simulation and Comparative Analysis of Drug Prioritization Algorithms in in-Vivo Trials

**DOI:** 10.1101/2025.08.08.669293

**Authors:** Ohad Landau, Kartheeswaran Thangathurai, Shai Magidi, Angel Porgador, Eitan Rubin

## Abstract

A critical challenge in personalized medicine is identifying the optimal drug treatment for individual patients based on their unique biological profiles. While recent advancements have led to a surge in drug prioritization algorithms utilizing patient omics data, including regression, classification, multiple kernel learning, deep learning, and AI-based methods, objective validation remains essential to discern their effectiveness. This study introduces” OptiRanker,” an innovative statistical framework designed to simulate and optimize the in vivo validation of these drug prioritization algorithms by ranking their predictive accuracy across varied conditions. OptiRanker not only mimics algorithmic predictions but also integrates them into ranked datasets, enabling the optimal selection of test subjects and drugs for trial setups. By providing a rigorous evaluation subset for ranking and comparison of high- and low-performing algorithms, OptiRanker facilitates the design of cost-effective in vivo trials tailored to efficiently assess predictive capability. This addresses a key bottleneck in personalized medicine: achieving a robust, scalable validation of computational models that can accelerate the clinical translation of promising algorithms. The efficient ranking provided by OptiRanker has the potential to advance personalized treatment strategies, ultimately leading to more effective, individualized patient care. The Python code for OptiRanker is available at the following GitHub repository: https://github.com/OhadLandau/OptiRanker.

## 1 Introduction

The field of precision oncology has seen remarkable advancements in recent years, with the development of increasingly sophisticated diagnostic tools and targeted therapies tailored to the unique molecular profiles of individual tumours [1]. This paradigm shift has been driven by a deeper understanding of the complex genomic and immunological landscapes of cancer, which has enabled the development of therapies that target specific molecular alterations or other biological characteristics implicated in disease initiation and progression [2–4] yet the process of prioritizing and evaluating potential therapeutic candidates remains a significant challenge. [5]

The rapid expansion of our knowledge about the molecular underpinnings of cancer has also revealed the inherent complexity and heterogeneity of these diseases, underscoring the need for individualized, gene-directed treatment approaches that go beyond the traditional tumor type-centric model [3]. In this context, the development of robust and reliable drug prioritization algorithms has emerged as a promising component of precision oncology, as clinicians and researchers seek to identify the most effective therapeutic strategies for each patient based on their unique characteristics [6]. However, the evaluation of these drug prioritization algorithms has been hampered by the inherent challenges of clinical trials, which often rely on small patient cohorts and may not capture the full spectrum of disease heterogeneity [2] or the high costs of such trials [7] and are further complicated by ethical considerations [8]. A further limitation in current clinical trial designs is the inability to test multiple drugs simultaneously on the same patient. This constraint is particularly problematic when various drug prioritization algorithms disagree on which therapeutic approach will elicit the best response. In such cases, only one drug can typically be tested in the patient at a time, preventing direct comparison and annotation of drug efficacy on a patient-specific basis. This limitation is especially relevant in the context of personalized medicine and when leveraging omics data.

The field of drug prioritization algorithms has made significant strides in recent years. A wide range of computational approaches have been developed that leverage various types of data, including genomic, transcriptomic, and clinical information. Each method has its own strengths and limitations [9]. For example, some algorithms have been designed to prioritize drugs based on their ability to target specific genetic alterations or molecular pathways implicated in cancer, [10,11] while others have leveraged machine learning techniques to identify patterns in large-scale datasets that may inform treatment decisions [12–21]. Critically, the performance of these algorithms can vary significantly based on the data and methods used, as well as the clinical context in which they are applied [2]. Furthermore, performance metrics evaluated on diverse datasets may introduce biases and may not accurately reflect real-world clinical scenarios [22] or act as comparable benchmarks [23]. Although retrospective analyses and early-phase clinical trials have shown promise in matching targeted therapies with specific genomic aberrations, some researchers have argued that the use of personalized treatment approaches needs to be further validated before being ready for widespread clinical implementation [24]. As a result, there remains a pressing need to rigorously validate the performance of these algorithms and optimize the *in vivo* trials to evaluate their efficacy in real-world clinical settings [7].

A key challenge in evaluating drug prioritization algorithms lies in the difficulty of establishing a reliable ground truth for comparison [25]. *In silico* validation, while valuable, often relies on datasets that may not fully represent the complexities of real-world patient populations and treatment responses [26]. Retrospective analyses of clinical trials can be confounded by factors such as patient selection bias and variations in treatment protocols [27]. Prospective clinical trials specifically designed to validate these algorithms are ideal but can be prohibitively expensive and time consuming, particularly given the need to evaluate multiple algorithms and their performance across diverse cancer types and patient subgroups.

The alternative to clinical trials is experimental *in vivo* models such as PDX mice, in which the barriers of clinical trials are much reduced or eliminated. It is possible, for example, to imagine a trial in which the same tumor is used to create multiple PDX mice and each is treated with the drugs selected by a different algorithm. Comparing the predicted efficacy to the observed response will help rank the prediction algorithms. However, with over 150 approved targeted drugs, and over 11,000 possible pairwise combinations, testing the performance of all is impractical. It is essential to choose *a priori* a smaller number of recommendations to test that will achieve a satisfactory ranking.

To this end, we have developed “OptiRanker,” a simulation-based framework that aims to generate the ideal combination of drugs and patient numbers to determine the best-to-worst order of prioritization algorithms in a time- and cost-effective approach. The core premise of the OptiRanker is to provide a platform where a quantitative approach to assess the potential performance of drug prioritization algorithms can be done, allowing researchers to test the robustness and accuracy of these algorithms under a variety of conditions and to identify the optimal experimental design for *in vivo* validation studies. By simulating the performance of different algorithms researchers can identify the most informative and efficient experimental setups. This approach has the potential to significantly reduce the time and cost associated with ranking drug prioritization algorithms in preclinical and clinical trials, ultimately accelerating the translation of these powerful tools to clinical practice.

## 2 Methods

### 2.1 Data Sources

#### 2.1.1 Simulated Data

##### Initialization of Ground Truth

A ground truth dataset was generated as a matrix of random permutations of integers ranging from 1 to the maximal number of drugs for each individual. This matrix represents the ideal or “ground truth” ranking of drugs for each individual (i.e., the outcome of a perfect drug prioritization algorithm).

##### Generating a series of Predictors with increasing Noise

Each predictor was generated by copying the ground truth dataset and introducing increasing levels of noise. The noise was introduced by randomly selecting individuals and performing a specified number of swaps within each individual’s drug rankings. For a given predictor *P*_*p*_:

- *p* individuals were randomly selected.
- For each selected individual, the rank of *p* random pairs of drugs was randomly swapped.

This process was repeated for every individual, resulting in predictors with decreasing accuracy as the predictor index *p* increased. A new 2D matrix of (*drugs* × *individuals*) was created for each predictor.

#### 2.1.2 Transcriptions

Expression profiles from the Cancer Cell Line Encyclopedia (CCLE)[28] and drug response metrics (IC50, GDSC)[29] for a total of 386 drugs and 1405 cell lines were obtained. Data was downloaded from the CCLE website (https://sites.broadinstitute.org/ccle/datasets, accessed on 16/11/2024 for both). Drugs and cell lines were filtered based on the following criteria:

- The drug was tested on all the chosen cell lines, therefore having experimentally determined IC50 values for all.
- The drug had data compatible with the selected algorithms; For example, Regorafenib—approved for metastatic colorectal cancer—was present in the WINTHER knowledge base but absent from DDPP.

These criteria ensured a large number of cell lines and drugs were chosen while ensuring all have IC50 values. The result is a dataset of 37 drugs and 570 cell lines, which were used in the in-silico experiments (Fig. 5).

### 2.2 DDPP, SIMS and WINTHER

Three algorithms, developed in part by the authors, were chosen: DDPP[30], SIMS[11], and WINTHER[10], Each was used to prioritize drugs based on transcriptomic data. These algorithms integrate multi-omic data to personalize cancer treatment strategies, each utilizing different combinations of genomic, transcriptomic, and other molecular data to generate predictions. A complete description of these algorithms is beyond the scope of this paper and all a described elsewhere [10,11,30]. Briefly, WINTHER measures the match between transcriptomic data and literature on drug efficacy. SIMS focuses on identifying disturbed intervention points through multi-omic data (e.g., genomic sequencing, copy number variation, transcriptomics, and miRNA expression). It elucidates activation patterns of 24 cancer-related “interventional nodes” and recommends single or combination therapies based on a scoring system that ranks activated nodes for each patient. DDPP aims to predict the change in predicted survival duration for each targeted therapy by analyzing gene expression profiles associated with drug sensitivity pathways. It utilizes a comprehensive, tumor- and treatment-agnostic approach, leveraging transcriptomic data to generate continuous survival estimates rather than binary response predictions. All three algorithms require expression profiles from matched normal tissues.

### 2.3 Scoring a Predictor

Each candidate predictor P_p_ is scored against the ground-truth by a position-biased weighted mean-squared error (WMSE)[31]. Briefly, squared rank deviations of a given predictor form the ground truth are aggregated across all individuals and drugs. The error is weighted by a coefficient (β) so that mistakes involving highly ranked drugs incur a steeper penalty. Afterwards, aggregated raw errors are inverted and linearly rescaled to the unit interval, yielding an intuitive score where larger values denote better agreement with the ground truth. In index notation, WMSE for a predictor P defined over I individuals and D drugs is

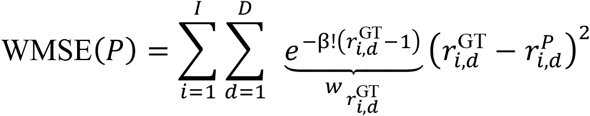

- 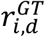 *and* 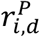 *are the ground truth and predicted ranks, respectively, for drug d in individual i*;
- 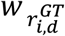 i*s the position weight assigned to that drug, controlled by the user selected discount factor β*

### 2.4 Sub-setting

To evaluate subsets of the data, the predictor rankings produced by these subsets were compared to rankings generated using the full dataset. The correlation between subset rankings and full-dataset rankings was calculated using the Spearman correlation coefficient [32]. In index notation, the standard Spearman correlation for two sets of ranks *x* and *y* of length *M* is:

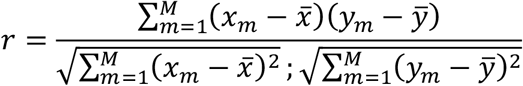

where *x*_*m*_ and *y*_*m*_ represent the corresponding rank values for the *m*-th element (e.g., drug or individual) in the subset and the full set, respectively, and 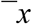 and 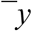 are their mean values.

#### Standard Deviation for Drug Selection

To identify the most informative drugs, the code sums individual-wise standard deviations (across predictors) for each drug:

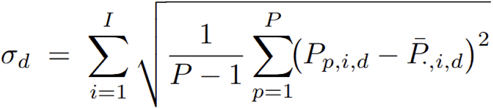

*Where* 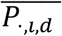 *denotes the average of P*_*p,i,d*_ *over all predictors. The d drugs with the largest* σ_*d*_ *values are then selected*.

#### Standard Deviation for Individual Selection

Similarly, for each individual *i*, the code computes per-drug standard deviations (across predictors) and sums them:

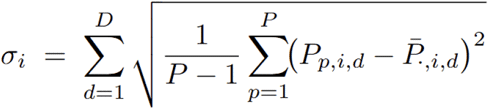

These selected drugs and individuals were then used to create subsets with the highest variability, ensuring the most informative data was utilized in further analysis (Fig. 2).

### 2.5 Cost-Effectiveness

To further improve OptiRanker’s utility, a cost-effectiveness analysis was carried out. Each subset whose correlation *ρ*_*i,d*_ (for a chosen number of individuals *i* and drugs *d*) exceeded a user given threshold (see below) was considered. The subset’s “cost” is simply the sum of individuals and drugs, *i* + *d*. The cost-effectiveness measure (in percentage) is computed as:

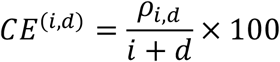

where *ρ*_*i,d*_ is set to zero if the correlation is below the threshold.

## 3 Results

### 3.1 OptiRanker Overview

As described in the methods section, we developed a method for simulating drug efficacy predictions with a known order of accuracy. We also describe how, given a ground truth, predictions can be ordered to capture their ability to predict drug ranking. The OptiRanker tool capitalizes on these capabilities to offer two modes of action (Fig. 1). In the fully simulated mode, users make critical decisions on the design of a trial, deciding how many drugs and how many samples would be required for any user-defined accuracy. This is achieved by creating simulated data in which “predictors” with decreasing accuracy are created by adding noise to the ground truth and are ranked using a score that estimates their accuracy. The correlation between the predictor rankings obtained with all the drugs and samples and the ranking calculated with fewer drugs and samples is then calculated for decreasing numbers of drugs and samples (see below). In the ‘Ranking’ mode, experimental results are used as ground truth, and actual prediction algorithms are ranked by their ability to reconstruct the ground truth.

**Figure 1:**
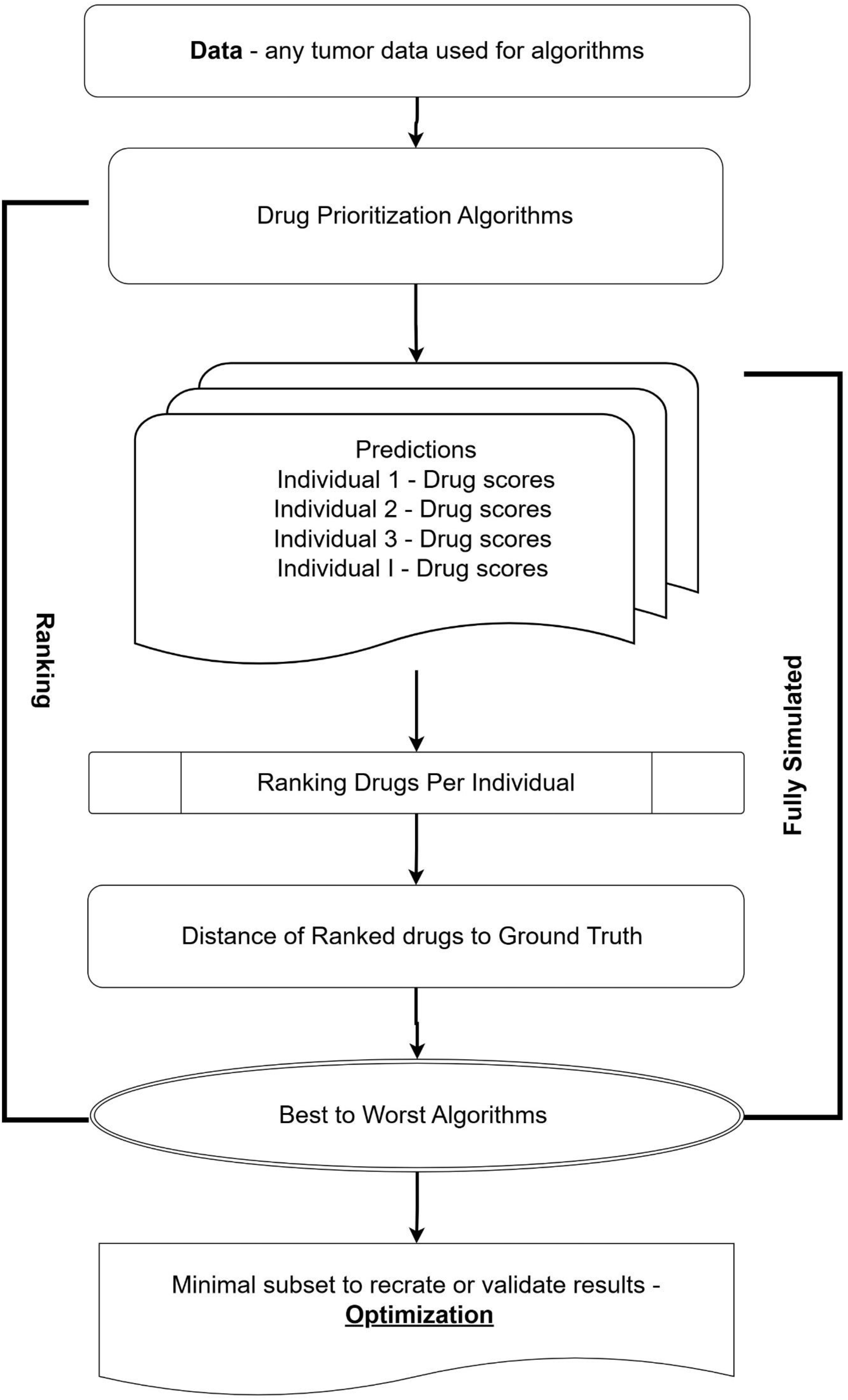
Workflow of OptiRanker. Generalized flow of simulating and optimizing Drug Prioritization Algorithms ranking experiments. Ground truth is either fully simulated or given by real world data, in our case – IC50 of drugs for cell lines, given by GDSC [29]

To test the quality of the simulation, we repeated it 100 times. For each generated dataset, an accuracy score was calculated for each predictor, and the results are summarized in Figure 2. As expected, predictors created with more noise receive lower scores, validating both our simulation and the score.

**Figure 2:**
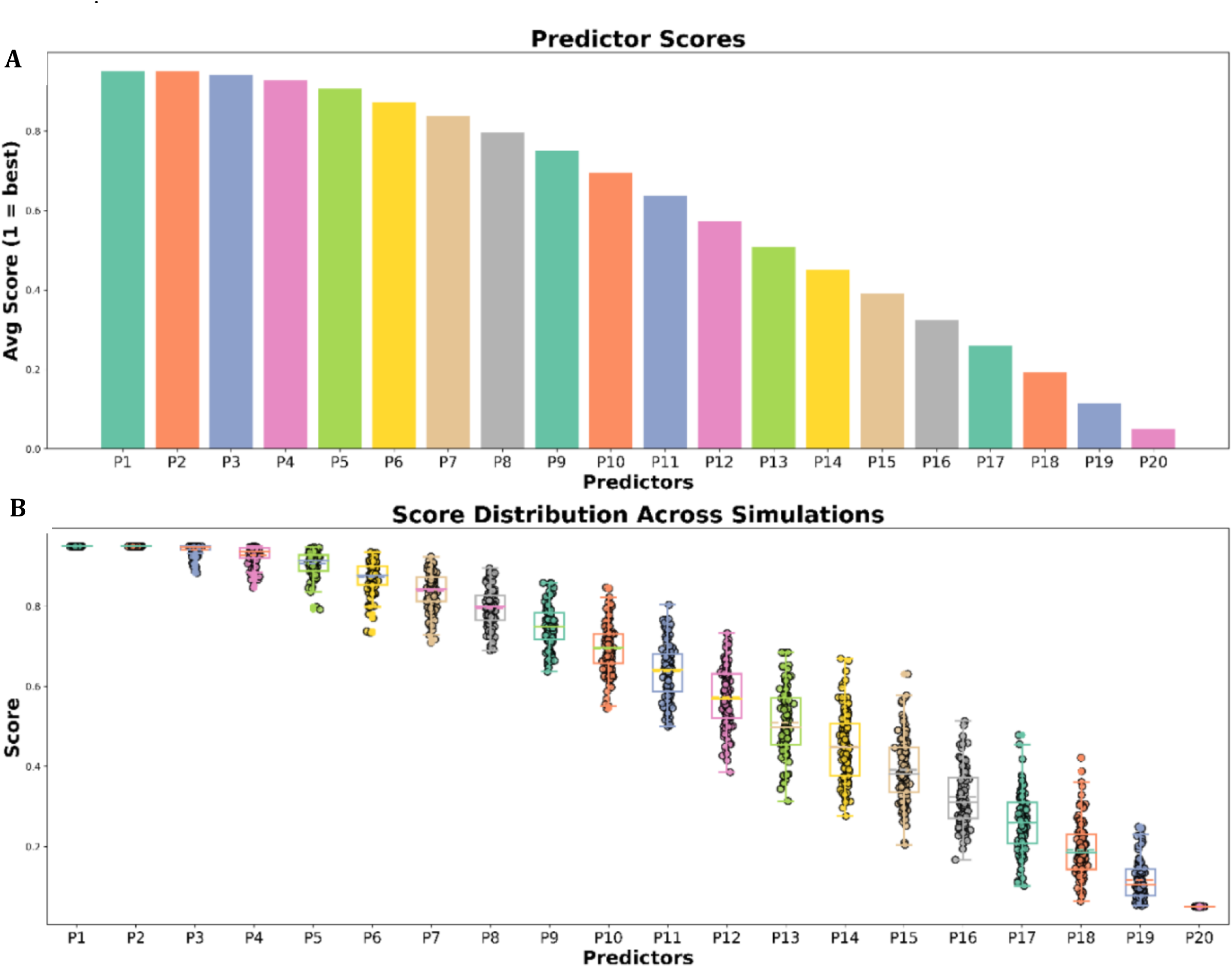
Data Generation and Noise Validation. A. Predictor scores were averaged across 100 simulations (individuals = 20, predictors = 20, drugs = 20), demonstrating the controlled noise induction and the ability of the simulation to correctly rank the predictors. B. Box plot of scores across all 100 simulations depicting the stochastic nature of the noise induction. Scores were MinMax scaled and inverted, as increased simulation size corresponded with greater noise and exponential scores.

**Figure 3:**
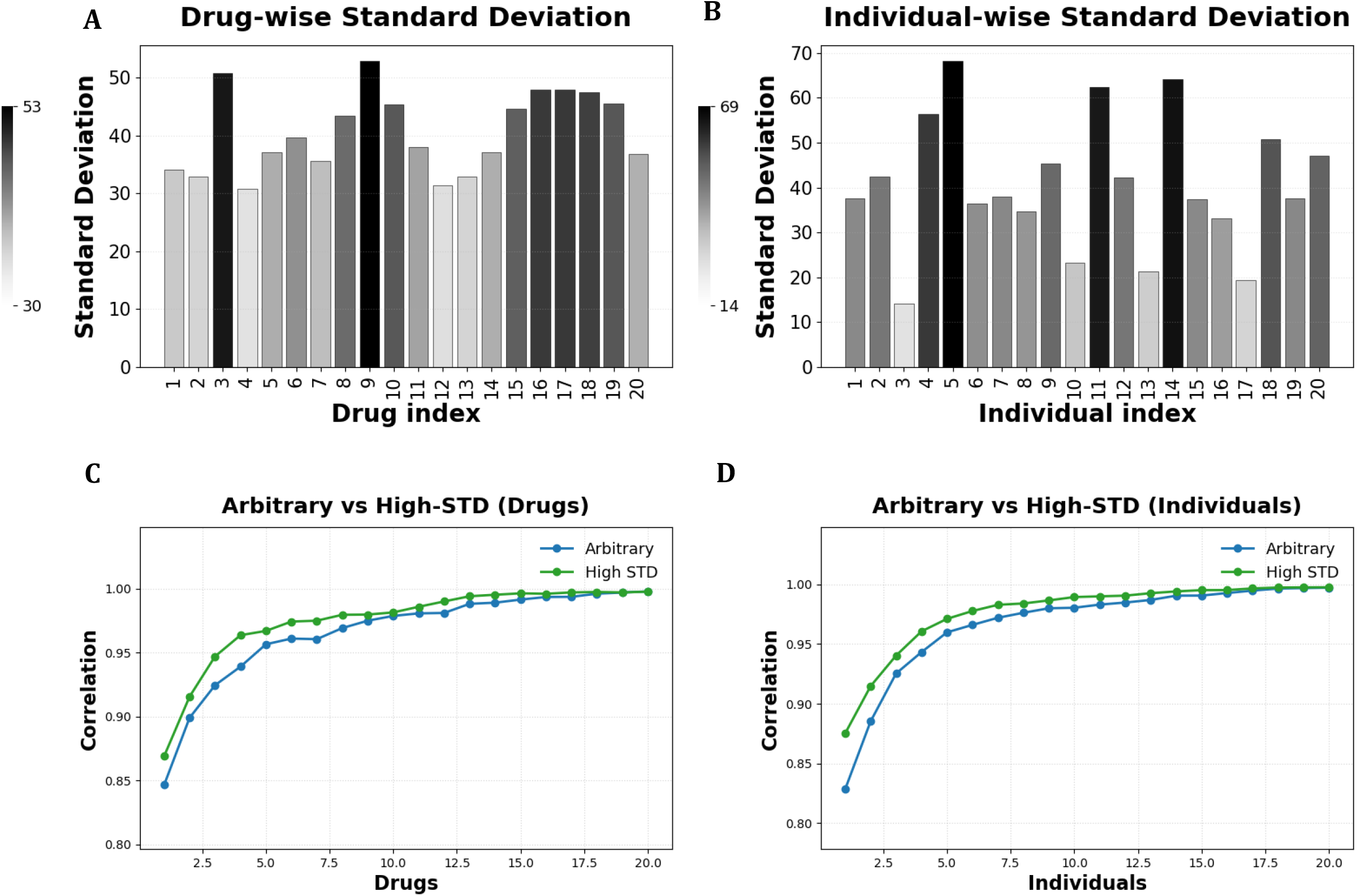
Informed Selection. A. Standard deviation of each drug, calculated across all individuals and all predictors. B. Standard deviation of each individual, calculated across all drugs and all predictors. Colours and height are proportional to standard deviation values. C, D. Predictor scores were recomputed on data subsets. In panel C all individuals were retained while the number of drugs was increased incrementally; in panel D all drugs were retained while the number of individuals was increased incrementally. For each subset size, two sampling strategies were compared: arbitrary selection and informed selection (based on the highest standard deviations in A or B). The resulting predictor rankings (best → worst) were Spearman-correlated with the ground-truth ranking. Simulations were performed with 100 iterations, 20 predictors, 20 drugs and 20 individuals, as described earlier.

### 3.2 Using Variance for Choosing Patients and Drugs

Next, we examined the value of informed selections of drugs and individuals. Our analysis (see below) requires subsetting the data by incrementally increasing the number of drugs and/or individuals from 1 to the limit provided by the user. Drugs and individuals can be randomly (“arbitrarily”) chosen or taken from a list sorted by standard variation. We calculated the standard deviation across all individuals for each drug, across predictions (Fig 2A) and, reciprocally, across all drugs for each individual (Fig 2B) to differentiate predictors (see 2.4) and accurately reconstruct the ranked order while relying on fewer observations than the full set. Spearman’s ρ climbed rapidly for both the informed drug-driven subsets (Fig 2C) and the informed individual-driven subsets (Fig 2D). The results highlight the importance of informed selection to the final score and ranked order, as evidenced by the significant immediate increase in the Spearman correlation coefficient, thus also illustrating its importance in subset optimization.

### 3.3 Trial Optimization

Trials can be optimized by choosing a subset of drugs and individuals to be tested. Fewer drugs and smaller trials require less resources but may not correctly rank the prediction algorithms. Using our simulated data, we can test the correlation obtained with all the drugs and individuals to the ranking obtained with subsets of drugs and individuals. The subsets of drug ranks within individuals for each simulated predictor were evaluated for their proximity to the ground truth results, and the correlation between the corresponding list of predictors ranked order and the known order when using all the dataset was then depicted (Fig. 4A). The strength of the correlation (r, Spearman correlation) between subsets increases as the increments become larger, the minimal strength specified by the user is highlighted to select the optimal subset for the investigator’s needs. As expected, predictor ranking was less accurate when fewer drugs and individuals were chosen. Nevertheless, very good accuracy (p>0.8) was observed with as few as 1, 2 or 3 individuals (with 5, 2 and 2 drugs, correspondingly). It should be noted here that these results are specific to the predictors we used, and a different number of drugs and individuals may be required to rank actual predictors. An interactive heatmap that allows the user to navigate through each subset and select the contents is then utilized for user-friendliness (Fig. S2). We then generated a cost-effectiveness heatmap illustrating how incremental increases in the subset size affect the prediction accuracy compared to the total number of individuals and drugs (Fig. 4B). The resulting trend highlights the potential for optimizing *in vivo* trials.

**Figure 4:**
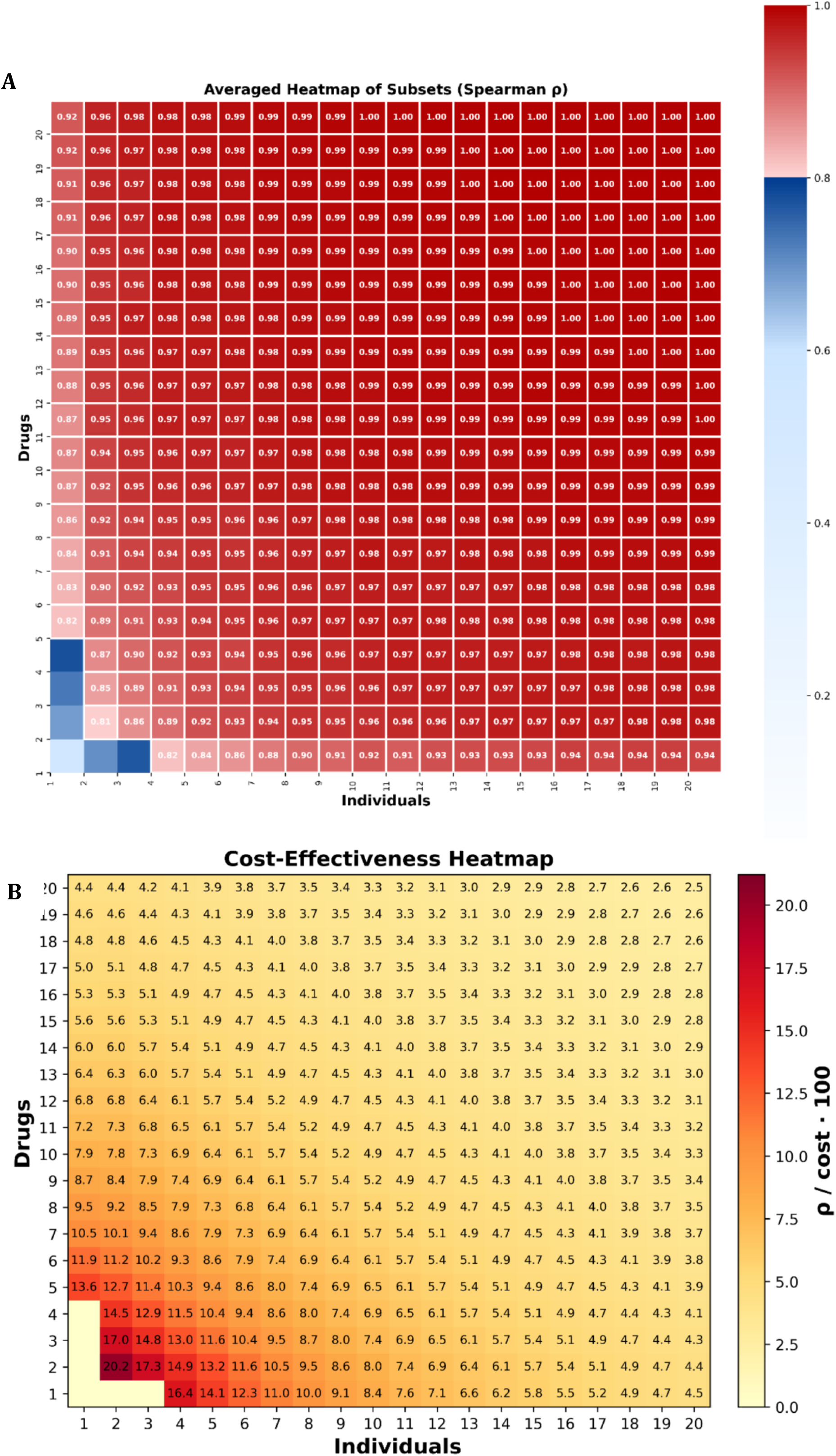
Heatmap of all possible subsets. A. Heatmap of the Spearman correlation coefficients between the ranked order of predictors generated from all possible informed subsets and the best-ranked order determined using the full dataset. The correlation values range from 0 (white), 0.5-0.8 (blue) and 0.8-1 (red), indicating the degree to which each subset can recapitulate the optimal ranking of predictors. Correlations above 0.8 (arbitrarily selected and user-defined) are highlighted. B. Cost-effectiveness heat map for all subset configurations that satisfy the specified correlation cut-off. Subsets whose correlation falls below the threshold are rendered uniformly in yellow, while those that meet or exceed it are colour-graded from light to deep red in proportion to their cost-effectiveness. Heatmap follows previous simulation structure: 20 drugs × 20 individuals × 20 predictors, repeated 100 times.

### 3.4 In-silico Trial

To test the ability of the simulation to propose how many drugs and individuals are necessary to rank prediction algorithms, we used cell line data from the CCLE trials. The trial that generated the data is described in detail elsewhere [28]. Briefly, it involved determining the IC50 of many drugs for many cell lines. Efficacy predictions of three in-house algorithms (DDPP [30], WINTHER [10], and SIMS [11]) were compared to measured IC50, suggesting the following order for the algorithms: DDPP, SIMS, WINTHER (Fig 5A). Median expression of every gene across selected cell lines was used as its normal value. Then, reducing predictions of the three algorithms and CCLE ground truth results into a 2D space using PCA (Fig 5B) resulted in cluster distances corresponding to the predictors ranked order obtained by OptiRanker. The same ranked order (DDPP, SIMS, WINTHER) was reconstructed using only one drug and six individuals (Supp Fig. S2). It is important to acknowledge that the right order of the three predictors may be achieved by chance. To address this, we tested this hypothesis by conducting 20 iterations of optimization using completely simulated data of the same dimensions – 3 predictors, 37 drugs and 570 individuals (i.e. cell lines). The integrity of the early minimal subsets was consistently recapitulated across iterations, adding strength and robustness to the optimized subset selection. This confirms the method we propose to reduce the number of drugs and individuals.

**Figure 5:**
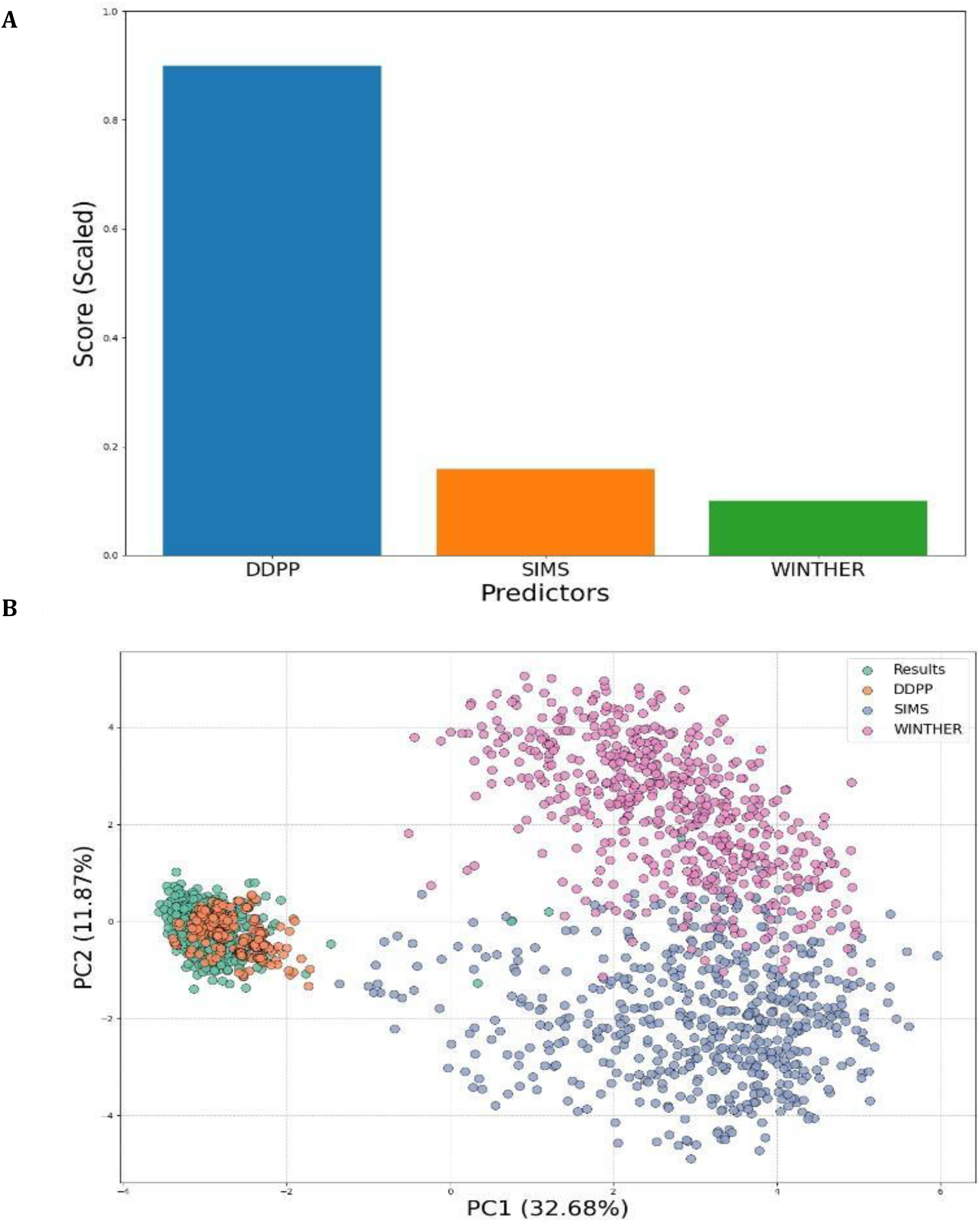
In-silico trial. Three algorithms—WIN [10], SIMS [11], and DDPP [30]—were applied to predict drug responses using cell line transcriptomic data for 570 cell lines. Subsequently, 37 matching drugs were ranked based on their corresponding IC50 values for each. The drug response predictions generated by each algorithm were then compared to the ground truth IC50 results and ranked accordingly. A. Bar plots depicting algorithm scores. B. Predictions of the algorithms with the CCLE results clustered in a reduced dimensional space (PCA).

## 4 Discussion

In this work, we present a novel simulation for ranking and an optimization tool for pre-clinical trials of drug prioritization algorithms, utilizing minimal resources to correctly rank the best-to-worst predictor, which, to the best of our knowledge, is the first of its kind. By transforming algorithm predictions into ranks of drugs, we were able to simulate a sequence of noisy predictors against tested drug responses. Then, we showed that subsets of the data was sufficient to recapitulate the simulated and known rank of predictors, given by Spearman coefficient to estimate the recapitulation accuracy. Finally, predictions from three distinct algorithms - WIN, SIMS, and DDPP - were then ranked against known drug response data, and the minimal subset of tests sufficient to recreate such ranking was selected to establish the proof of concept.

The primary contribution of this work is the demonstration of the capability to objectively differentiate between drug prioritization algorithms, from strong to weak performers, by an experiment with optimal resource allocation and less then the entirety of data. We present “OptiRanker”, a framework that enables the design of efficient *in vivo* trials that will evaluate drug prioritization algorithms. A systematic and quantitative approach to evaluate the performance of drug prioritization algorithms is crucial for their future translation to clinical practice, and OptiRanker increases the feasibility of such an evaluation. Moreover, this approach is robust and can be used to assess seemingly different algorithms, often utilizing different modalities to prioritize drugs so long as they rank the same drugs. The validity of this framework is supported not only by extensive simulations exploring diverse predictions and drug rankings but also through *in silico* trial based on published algorithms and drug response databases. These validation steps demonstrate the reliability and applicability of the “OptiRanker” in real-world *in vivo* settings (even if not in clinical setting).

Although our methodology has demonstrated effective results, certain limitations have been identified that will be addressed in future works, if feasible. Primarily, the approach of utilizing rank-based predictions from various algorithms does not account for the magnitude of difference between ranks, which could potentially lead to penalizing minor differences or erroneously rewarding substantial changes. While Weighted Mean Square Errors (WMSE) scoring mitigates the potentially erroneous nature of value to rank transitions, exponentially penalizing ranks might not capture real-world magnitudes of positional distances. This issue remains elusive and requires further research. Furthermore, a significant challenge emerges as the number of algorithms, drug candidates, and participant cohorts increases. This escalation necessitates larger sample sizes to derive meaningful Spearman correlation coefficients. However, this requirement may lead to impractical or unrealistic trial designs. Conversely, reducing the number of predictors can result in extremely small subsets of participants and drug candidates, potentially yielding high correlations by mere chance rather than true significance. This problem of contrast is a critical consideration in any *in vivo* study and is not unique to OptiRanker: how one balances statistical power with practical feasibility?. Finally, additional validation and benchmarking efforts are required to fully demonstrate the advantages of the proposed framework, which can be achieved through expanding the *in silico* trial size and transitioning to *in vivo* validation studies.

Several studies have underscored the challenges associated with validating drug prioritization algorithms using traditional methods [33,34]. These challenges stem from the difficulty in establishing a reliable ground truth, the limitations of *in silico* datasets, and the high costs associated with prospective clinical trials. OptiRanker directly addresses the latter challenges by providing an efficient means to pre-clinically optimize *in vivo* trials. Future research will focus on addressing the limitations of the current framework, including identifying and incorporating a better and informed penalization of rank changes, exploring alternative correlation metrics for larger datasets, increasing in-silico trial size and integrating additional data modalities beyond transcriptomics. Moreover, collaborations with experimental groups will be actively pursued to validate the “OptiRanker” in real-world *in vivo* settings, further strengthening its translational potential. The development of such robust and efficient validation frameworks is essential for accelerating the deployment of drug prioritization algorithms.

## 5 Conclusion

This work introduces the “OptiRanker”. By leveraging simulations and rank-based analyses, our approach enables the identification of minimal yet informative experimental designs. While further validation and refinements are necessary, the “OptiRanker” represents a step towards the efficient and cost-effective evaluation of drug prioritization algorithms, ultimately contributing to the advancement of personalized medicine. This aligns with the growing recognition in the field that robust and generalizable methods for evaluating computational approaches in drug discovery are crucial [35]. Additionally, the utilization of *in silico* simulations to guide experimental design, as showcased in this study, is gaining increasing traction in various areas of biomedical research [36], highlighting the timeliness and relevance of this work.

## Code and Data Availability

All code and data used for the simulations, in-silico trial, scoring, and optimization processes presented in this paper are openly available on the GitHub below. This repository ensures reproducibility of the results and facilitates further research on the methodologies described. https://github.com/OhadLandau/OptiRanker

## Supplementary Information

**Supplementary S1.**
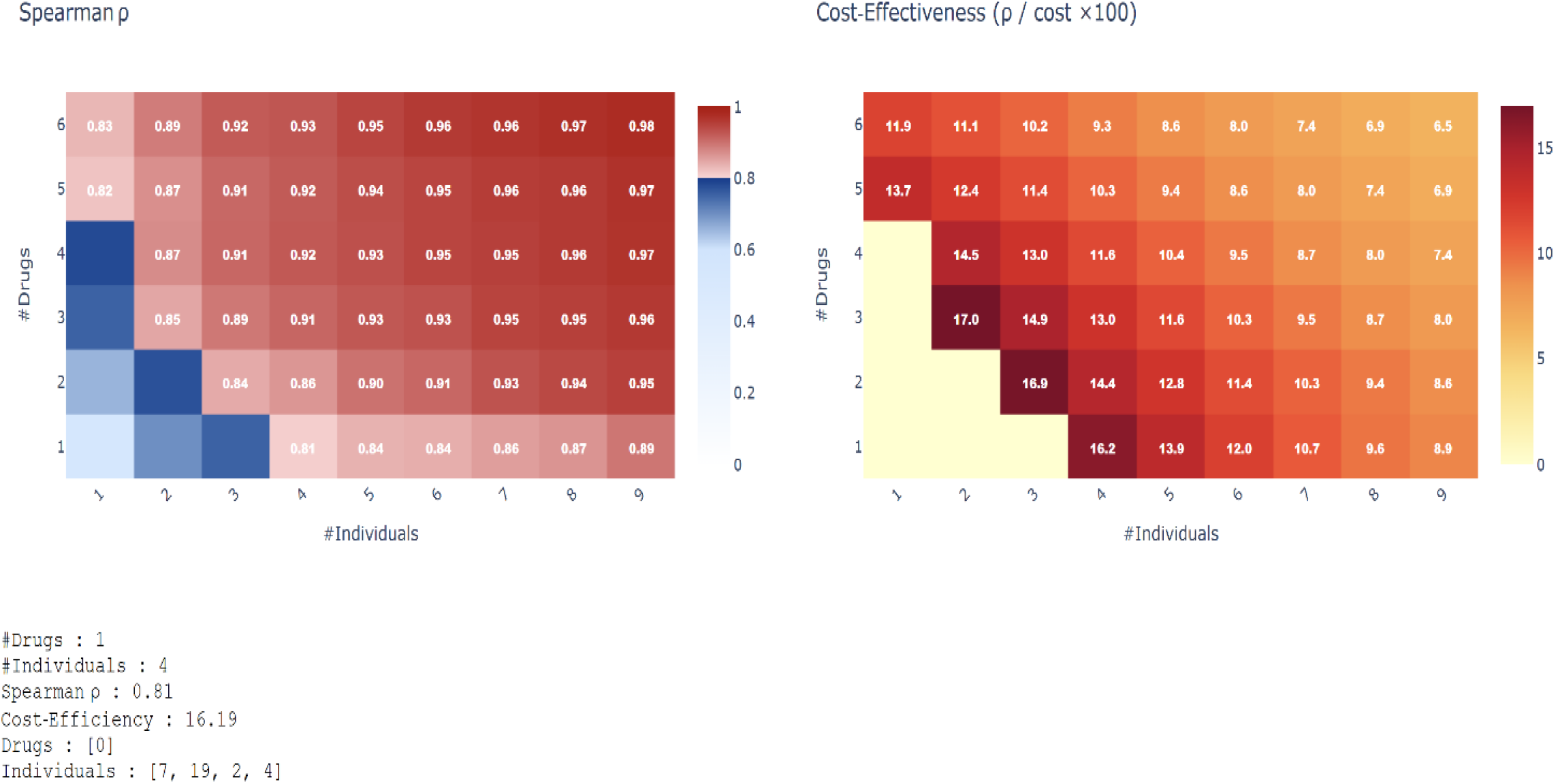
Snapshot from interactive heatmap generated by Plotly, displayed on local Dash app. Clicked cell displays the content of the selected subset. In this case, cell i4d1 with 0.81 correlation was selected.

**Supplementary S2.**
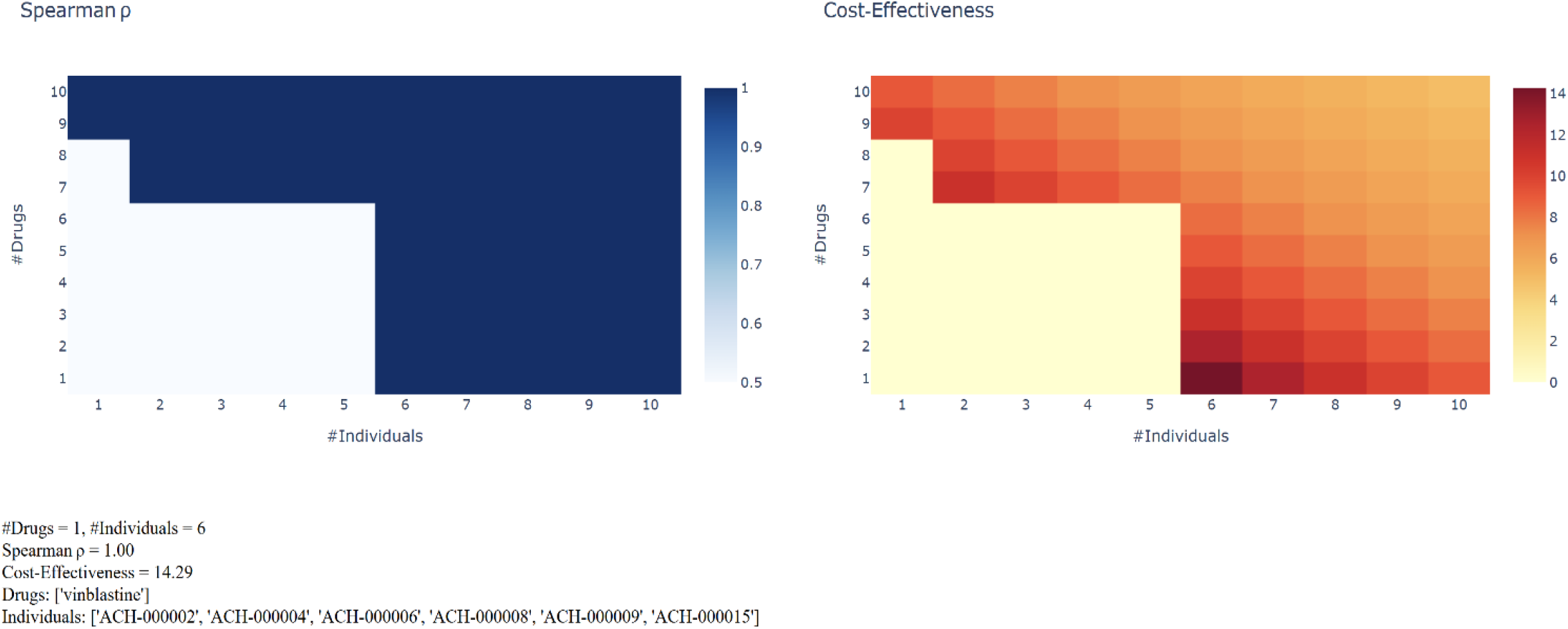
Snapshot of zoomed-in (10×10) interactive heatmap of In-Silico trial (full dimensions – 37 drugs X 570 cell lines X 3 predictors), with cost-effective analysis, generated by Plotly, displayed on local Dash app. Clicked cell displays the content of the selected subset. In this case, cell i6d1 with a correlation of 1 and cost-effectiveness of 14.29

